# Pluripotent stem cell-derived siTNK cells attack tumors via a synthetic CD8-CD28-TCR complex

**DOI:** 10.1101/2025.07.11.663392

**Authors:** Xiujuan Zheng, Fan Zhang, Yunqing Lin, Fangxiao Hu, Qitong Weng, Pengcheng Liu, Zhiqian Wang, Chenyuan Zhang, Yanhong Liu, Lijuan Liu, Yanping Zhu, Ziyun Xiao, Yao Wang, Leqiang Zhang, Hanmeng Qi, Yiyuan Shen, Yi Chen, Jiaxin Wu, Jiacheng Xu, Yaoqin Zhao, Tongjie Wang, Dehao Huang, Chengxiang Xia, Jinyong Wang, Mengyun Zhang

## Abstract

Tumor-associated antigen-specific T cell receptor (TCR)-engineered T cells offer a promising strategy for cancer therapy. Natural killer (NK) cells exhibit broad anti-tumor activity with low side effects but lack the capacity to recognize intracellular antigens. Here, we found that the human pluripotent stem cell (hPSC)-derived iNK cells, unlike tissue-isolated NK cells, expressed all four CD3 subunits at the transcriptome level. We introduced a synthetic gene-expressing complex (SCOTR), encoding a tumor antigen-specific TCR, CD8 coreceptor, and CD28 costimulatory molecule, into hPSCs to generate SCOTR-hPSCs. The SCOTR-hPSCs gave rise to abundant synthetic TCR complex-expressing iNK (siTNK) cells via an organoid induction method. These siTNK cells showed HLA-dependent, antigen-specific cytotoxicity against tumor cells and significantly suppressed tumor growth in tumor xenograft animal models, while also preserving universal non-specific tumor-killing activity. Collectively, siTNK cells, as hPSC-derived hybrid cells with dual features of adaptive T and inherent NK cells, offer an artificial cell source for human immunotherapies.

## Introduction

Immune cell therapy has been extensively applied in treating cancer patients^1^. In particular, chimeric antigen receptor T (CAR-T) cells, via targeting cell surface antigens, have demonstrated efficacy in treating B cell-related blood cancers^2,3^. However, most tumor-associated antigens (TAAs) are intracellular and presented on tumor cell surfaces as peptide-major histocompatibility complex (pMHC)^4^. T cells kill tumors mainly through TCR recognition of the pMHC complex, which is the main immunological mechanism of the body’s defense against tumor occurrence and progression^5^. TCRs recognizing specific TAAs are isolated and introduced into the T cells to generate TCR-T cells, which are employed to treat various types of cancers^6–8^. However, clinical translation of TCR-T cells for cancer therapy faces challenges, mainly because exogenous TCRs and endogenous TCRs within T cells are prone to mismatch, weakening TAA-TCR specificity and even creating off-target effects that cause autoimmune problems^9–11^. The risks including cytokine release syndrome (CRS), graft-versus-host disease (GVHD), immune effector cell-associated neurotoxicity syndrome (ICANS), and the inability to recognize tumor cells that downregulate human leukocyte antigen class I (HLA-I) molecules have also limited the application of T cell- based therapies^12,13^.

NK cells offer several advantages over T cells, including minimal toxicity, the ability to target abnormal cells without prior sensitization, and broad-spectrum tumor-killing characteristics^14,15^. Engineering NK cells with an antigen-specific CAR structure has demonstrated significant potential in treating malignant tumors^16,17^. However, neither NK cells nor CAR-NK cells possess the T cell-specific adaptive immune function to recognize tumor intracellular antigens. Overexpression of CD3 molecules (CD3δ, CD3γ, CD3ε, CD3ζ) and TCRs in NK cells demonstrated TCR-mediated specific anti-tumor activity^18–20^. Nevertheless, NK cells derived from human tissues face challenges such as high heterogeneity, low efficiency of multi-gene engineering, and high costs^21^. Furthermore, NK cell lines, such as NK-92, must be irradiated before infusion to prevent tumorigenesis, which may impact the overall therapeutic potential^22^. Human pluripotent stem cells (hPSCs), including human embryonic stem cells (hESCs) and induced pluripotent stem cells (iPSCs), offer advantages such as unlimited cell sources and clonal screening convenience for multiple genetic modifications^23–25^. They are promising cell sources for producing standardized, off-the-shelf engineered NK cells.

During the process of TCR signal transduction in cytotoxic T lymphocytes, CD8 and CD28 play important roles. CD8, existing either as an αα homodimer or an αβ heterodimer, acts as a coreceptor to enhance the stability of the TCR-pMHC interaction by binding to MHC class I molecules^26,27^. CD28, as a costimulatory molecule, can strengthen the TCR signal (the first signal) by interacting with CD80 or CD86 to transmit the essential costimulatory signal (the second signal)^28,29^. Introducing TCR, CD3, CD8, and CD28 molecules simultaneously into NK cells may endow NK cells with superior adaptive immune function.

In this study, hPSC-derived iNK cells, unlike tissue-isolated NK cells, were confirmed to express all four subunits of CD3 (*CD3D*, *CD3E*, *CD3G*, and *CD247*) at the transcriptome level. We subsequently introduced the TCR, CD8 coreceptor, and CD28 costimulatory molecule gene-expressing elements into hPSCs to construct SCOTR- hPSCs, including SCOTR-hESCs and SCOTR-iPSCs. These SCOTR-hPSCs successfully gave rise to abundant synthetic TCR complex-expressing iNK (siTNK) cells. Serial functional assays indicated that the siTNK cells possessed superior intracellular antigen-targeting cytotoxicity compared with hPSC-derived iNK cells. Notably, the siTNK cells also exhibited significantly greater cytotoxicity than conventional TCR-T cells. Taken together, we have established artificial siTNK cells from engineered hPSCs, which maintain the inherent immune characteristics of NK cells and acquire the pMHC-TCR-mediated specific cytotoxic capability exclusive to T cells. siTNK cells provide a brand-new cell resource for cancer treatment with significant public health implications.

## Results

### SCOTR complex engineering of hPSCs and derivation of induced EBV-siTNK cells

In our previous study, we successfully generated a large number of iNK cells through the differentiation of hPSCs^23^. iNK cells, similar to primary NK cells, showed low expression of *CD8A* and lacked the expression of *CD8B* and *CD28* at the transcriptome level. Intriguingly, iNK cells expressed all four subunits of CD3, which differed from primary NK cells (**Fig. 1a,b**). Given that iNK cells expressed the complete CD3 subunits, we hypothesized that the exogenous TCR could be efficiently assembled into a TCR-CD3 complex when introduced into iNK cells. To test this hypothesis, we used retroviral infection to introduce an Epstein-Barr virus (EBV)-specific TCR^30^, which recognizes the epitope of EBV-LMP2 protein (SSCSSCPLSK) presented by HLA- A*11:01, into both iNK and primary NK cells to assess the TCR-CD3 complex assembly. The TCRα chain, consisting of TRAV22, TRAJ43, and TRAC, and the TCRβ chain, consisting of TRBV4-1, TRBD1, TRBJ2-7, and TRBC2, along with the GFP reporter, were linked successively by the 2A peptide sequence (**Fig. 1c**). As anticipated, 72.7% of GFP^+^ iNK cells successfully assembled the TCR-CD3 complex, in contrast to just 9.7% in primary NK cells (**Fig. 1d**). To efficiently generate TCR-engineered iNK cells with high homogeneity and low cost, we introduced *piggyBac* (PB) plasmids containing exogenous TCR-expressing elements into hESCs. Given that TCR expression cannot be detected at hESCs due to the absence of CD3, we used GFP as a reporter. To enhance TCR affinity and strengthen the signal transduction of the TCR complex, we simultaneously introduced CD8 and CD28 expression elements into hESCs (**Fig. 1e**). After two rounds of sorting, these genetically engineered hESCs were defined as EBV SCOTR-hESCs (**Fig. 1f**). Flow cytometry confirmed the expression of these exogenous components in EBV SCOTR-hESCs. Approximately 98% of EBV SCOTR-hESCs expressed EBV TCR-GFP, CD8, and CD28 (**Fig. 1g**). These modified hESCs were subsequently differentiated into EBV-siTNK cells via an efficient organoid induction method (**Fig. 1h**). The resulting EBV-siTNK cells expressed high levels of CD45, CD56, and EBV TCR (**Fig. 1i**). In addition, the EBV-siTNK cells simultaneously expressed CD8α, CD8β, CD28, and CD3 (**Fig. 1j**). EBV-siTNK cells displayed similar characteristics to iNK cells, including the expression of NK cell activation and inhibitory receptors, effector molecules, and non-specific cytotoxic activity against tumor cells **(Extended Data Fig. 1a,b)**. These results suggest that we have established an efficient method for generating siTNK cells through hPSC engineering and induction.

**Fig. 1.**
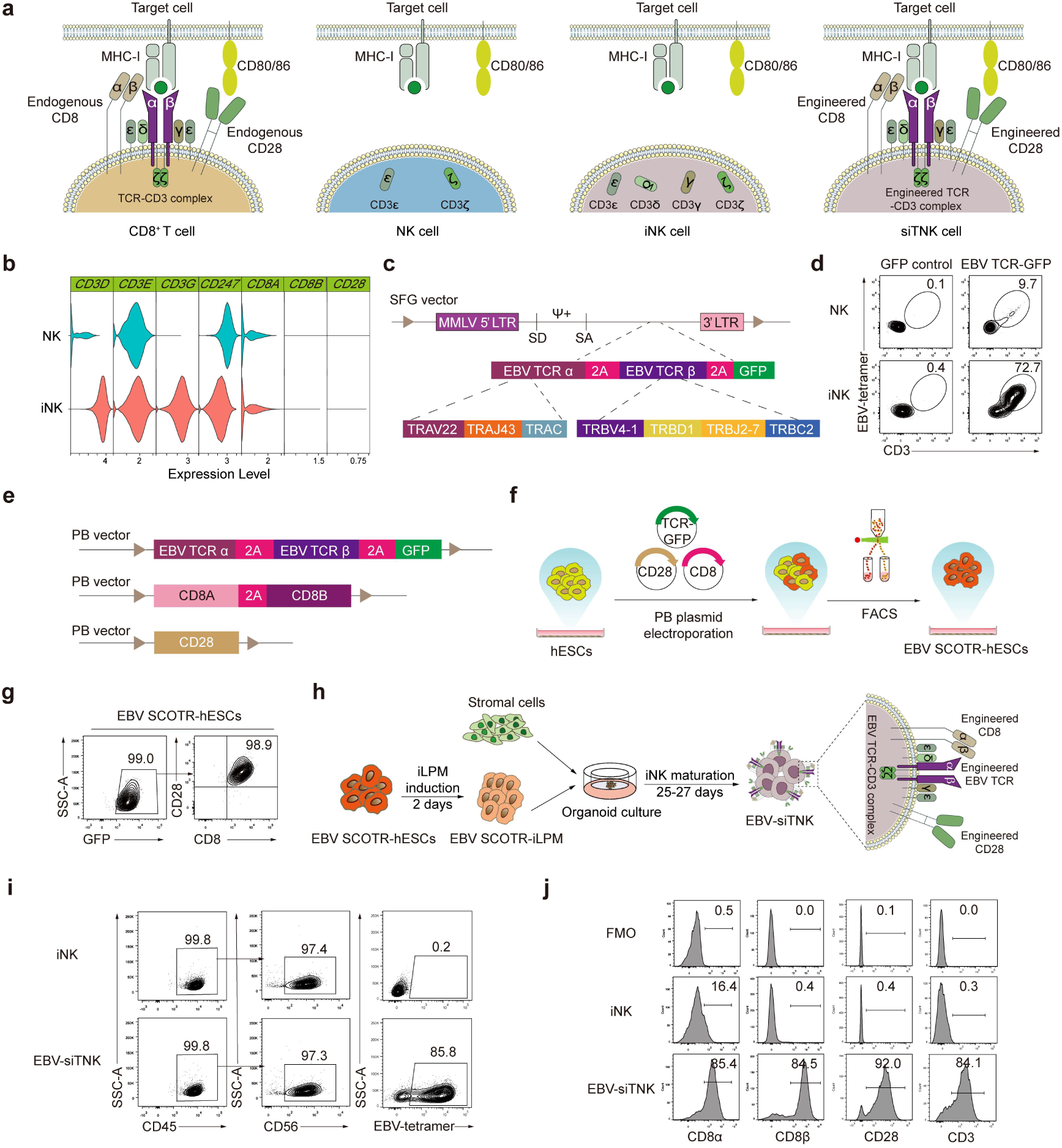
Engineering of hESCs and derivation of induced EBV-siTNK cells. **a**, The schematic structures of the molecules associated with the T-cell receptor (TCR) are shown for different cell types: CD8^+^ T cells, umbilical cord blood (UCB)-derived NK cells, hPSC-derived iNK cells, and predicted siTNK cells. The TCR-CD3 complex consists of two main TCR polypeptide chains, TCRα and TCRβ, along with four CD3 chains: CD3γ, CD3δ, CD3ε, and CD3ζ. CD8: Coreceptor. CD28 and CD80/86: Costimulatory molecules. **b**, Transcriptome characterization of CD3 subunits (*CD3D*, *CD3E*, *CD3G*, *CD247*), CD8 subunits (*CD8A*, *CD8B*), and CD28 (*CD28*) expressed in NK and iNK cells. NK cells were isolated from UCB and expanded *in vitro* for 6 days. iNK cells were differentiated from hESCs, which were collected on day 27 of induction. The raw data of NK and iNK cells were downloaded from GSA (HRA007978, HRA001609), and the gene expression levels were presented using violin plots. **c**, Schematic of the design of SFG-EBV TCR-GFP retroviral vector. The α and β sequences of the EBV TCR and GFP reporter were linked successively by the 2A peptide sequence. The α chain consisted of TRAV22, TRAJ43, and TRAC. The β chain consisted of TRBV4-1, TRBD1, TRBJ2-7, and TRBC2. MMLV, Moloney murine leukemia virus. LTR, long terminal repeat. SD, splice donor. ψ+, packaging signal. SA, splice acceptor. **d**, EBV TCR (detected by EBV-specific tetramer) and CD3 (detected by CD3ε antibody) expression levels in transfected NK or iNK cells (gated on CD45^+^CD56^+^GFP^+^ cells). **e**, The design of the PB vector constructs for the EBV TCR, CD8 coreceptor, and CD28 costimulatory molecule. **f**, Strategy diagram for the construction of EBV SCOTR-hESCs. **g**, Flow cytometry analysis of EBV TCR-GFP, CD8 (detected by CD8α antibody), and CD28 expression levels in EBV SCOTR-hESCs. **h**, Schematic diagram of the induction of EBV-siTNK cells. iLPM: induced lateral plate mesoderm cells. **i**, Representative flow cytometry plots of iNK and EBV-siTNK (CD45^+^CD56^+^EBV TCR^+^) cells. **j**, Representative histogram plots of CD8α, CD8β, CD28, and CD3 expression levels in iNK and EBV-siTNK cells.

### EBV-siTNK cells exhibit HLA-dependent, intracellular antigen-specific cytotoxicity against tumor cells *in vitro*

To assess the antigen-specific killing activity of EBV-siTNK cells, we conducted a cytotoxicity assay using the CD80^+^CD86^+^ HMy2.CIR tumor cell line (**Extended Data Fig. 2a**), which was transfected with HLA-A*11:01 (C1R-1101 cells). Meanwhile, we generated EBV TCR-T cells, which were used to evaluate whether EBV-siTNK cells possess superior tumor-killing activity compared with EBV TCR-T cells (**Fig. 2a**). The EBV TCR-positive T cells were sorted for *in vitro* killing assay. C1R-1101 cells were pre-incubated with the EBV-LMP2 peptide (SSCSSCPLSK) for 20 h and then co-cultured with iNK, EBV TCR-T, or EBV-siTNK cells for 4 h (**Fig. 2b**). The results showed that the cytotoxicity of EBV-siTNK cells against EBV peptide-pulsed C1R- 1101 cells was significantly higher than that of iNK cells, indicating that equipping iNK cells with TCR enhanced their tumor-targeted cytotoxicity in the presence of the coreceptor CD8 and the costimulatory molecule CD28. Notably, compared with EBV TCR-T cells, EBV-siTNK cells exhibited superior cytotoxic activity (**Fig. 2c**). To simulate the intracellular antigen presentation process by tumors, HLA-A*11:01 and the EBV-LMP2 peptide-expressing elements were introduced into CD80^+^CD86^+^ Raji cells (Raji-1101-EBV peptide cells) (**Extended Data Fig. 2b**), which were then co- incubated with iNK or EBV-siTNK cells. As expected, EBV-siTNK cells exhibited significantly higher cytotoxicity (**Fig. 2d,e**). To further assess the persistent cytotoxicity of EBV-siTNK cells, we conducted three rounds of tumor-killing assays with Raji- 1101-EBV peptide cells (**Fig. 2f**). The results indicated that EBV-siTNK cells exhibited a robust serial tumor-killing activity that was superior to that of iNK cells (**Fig. 2g**). Moreover, EBV-siTNK cells demonstrated higher expression levels of CD107a, TNF- α, and IFN-γ when stimulated with Raji-1101-EBV peptide cells (**Fig. 2h-m**).

**Fig. 2.**
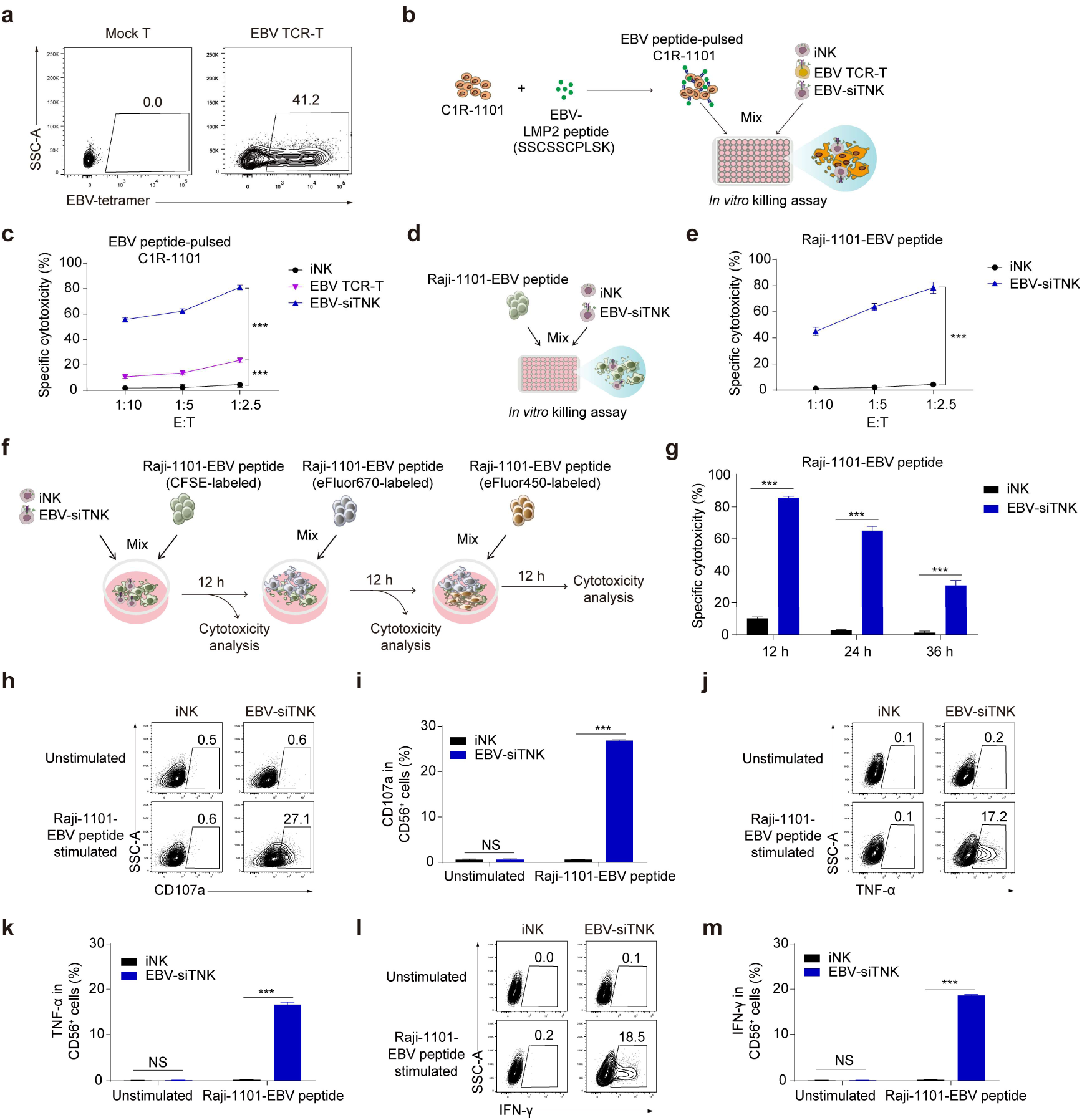
Evaluation of the tumor-killing activities of EBV-siTNK cells *in vitro*. **a**, Flow cytometry analysis of EBV TCR expression levels in Mock T (unmodified) and EBV TCR-T (infected with EBV TCR retroviruses) cells. **b**, Schematic diagram of measuring the cytotoxicity of iNK, EBV TCR-T, and EBV-siTNK cells against C1R- 1101 antigen-presenting cells exogenously pulsed with EBV-LMP2 peptide (SSCSSCPLSK). After pre-incubating C1R-1101 cells with the peptide (1 μM) for 20 h, the C1R-1101 cells were co-cultured with iNK, EBV TCR-T, and EBV-siTNK cells for 4 h, respectively. **c**, Cytotoxicity analysis of iNK, EBV TCR-T, and EBV-siTNK cells against C1R-1101 cells (1 × 10^4^) pulsed with EBV peptide at the indicated effector to target (E:T) ratios (*n* = 4 per group). **d**, Schematic diagram of measuring the cytotoxicity of iNK and EBV-siTNK cells against Raji-1101-EBV peptide cells. The Raji-1101-EBV peptide cells were co-cultured with iNK or EBV-siTNK cells for 6 h. **e**, Cytotoxicity analysis of iNK and EBV-siTNK cells against Raji-1101-EBV peptide cells (1 × 10^4^) at the indicated E:T ratios (*n* = 4 per group). **f**, Experimental design for the serial killing assays of iNK and EBV-siTNK cells. CFSE-labeled Raji-1101-EBV peptide cells (1 × 10^4^) were co-cultured with iNK or EBV-siTNK cells at the E:T ratio of 1:1. After 12 h, fresh eFluor670-labeled Raji-1101-EBV peptide cells (1 × 10^4^) were added and co-cultured for another 12 h. At 24 h, fresh eFluor450-labeled Raji-1101- EBV peptide cells (1 × 10^4^) were added and co-cultured for an additional 12 h. The death rates of CFSE^+^ cells (12 h), eFluor670^+^ cells (24 h), and eFluor450^+^ cells (36 h) represented the cytotoxicity of iNK or EBV-siTNK cells. **g**, Statistics of specific cytotoxicity of iNK and EBV-siTNK cells in serial killing assays (*n* = 4 per group). **h**- **m**, Flow cytometry analysis and statistical analysis of the expression levels of CD107a (**h**, **i)**, TNF-α (**j**, **k**), and IFN-γ (**l**, **m**) in iNK and EBV-siTNK cells which were stimulated with Raji-1101-EBV peptide cells or not (E:T = 1:2) for 4 h (*n* = 3 per group). Data were presented as mean ± SD. Statistics: two-tailed independent *t*-test (**c**, **e**, **g**, **i**, **k**, **m**) or Mann-Whitney U test (**m**). NS, not significant, \*\*\**p* < 0.001.

The iPSC-derived immune cell therapies offer several benefits, including the potential for patient-specific autologous treatments that minimize immune rejection, as well as allogeneic “off-the-shelf” options derived from HLA-matched donors for broader applicability. To verify whether the SCOTR complex engineering strategy is applicable to iPSCs, we introduced PB plasmids containing CD8, CD28, and EBV TCR-GFP coding sequences into iPSCs and established the EBV SCOTR-iPSCs **(Extended Data Fig. 3a)**. Subsequently, through *in vitro* induction and differentiation, we successfully obtained iPSC-EBV-siTNK cells **(Extended Data Fig. 3b,c)**. Further *in vitro* functional assays revealed that iPSC-EBV-siTNK cells exhibited superior target-killing activity compared with iPSC-iNK cells **(Extended Data Fig. 3d,e)**. These results confirmed that the SCOTR complex empowered iNK cells with HLA-dependent, intracellular antigen-specific, and enhanced cytotoxicity *in vitro* by mimicking TCR signaling of CD8^+^ T cells.

### WT1-SCOTR empowers iNK cells with intracellular antigen-specific cytotoxicity against tumor cells

To determine whether the SCOTR complex expression strategy in iNK cells is applicable to general TCRs, we further tested the WT1 TCR, which recognizes the epitope of WT1 protein (RMFPNAPYL) presented by HLA-A*02:01 and has been extensively studied in tumor killing. We first introduced the PB vectors containing WT1 TCR-GFP, CD8, and CD28 elements into hESCs to establish the WT1 SCOTR-hESCs (**Fig. 3a,b**). Flow cytometry analysis confirmed the expression of the exogenous components in WT1 SCOTR-hESCs. Over 98% of WT1 SCOTR-hESCs expressed WT1 TCR-GFP, CD8, and CD28 (**Fig. 3c**). We subsequently differentiated these cells into WT1-siTNK cells (**Fig. 3d**). The resulting WT1-siTNK cells expressed high levels of CD45, CD56, CD16, and WT1 TCR (**Fig. 3e**). Additionally, these cells co-expressed high levels of CD8α, CD8β, CD28, and CD3 (**Fig. 3f**). To evaluate the antigen-specific killing activity of WT1-siTNK cells, we conducted a cytotoxicity assay using the HMy2.CIR cells transfected with HLA-A*0201 sequence (C1R-0201 cells) (**Fig. 3g**). The results showed that the WT1-siTNK cells exhibited superior cytotoxicity against WT1 peptide-pulsed C1R-0201 cells (**Fig. 3h**). We further used the WT1-expressing HLA-A*02^+^ BV173 cells that overexpress CD80 and CD86 to investigate the cytotoxicity of WT1-siTNK cells (**Fig. 3i**). Similarly, WT1-siTNK cells possessed TCR-mediated, CD8- and CD28-enhanced intracellular recognition of target antigens and cytotoxicity of target cells (**Fig. 3j,k**). Thus, the SCOTR complex strategy can be employed to evaluate the functionality of multiple TCRs.

**Fig. 3.**
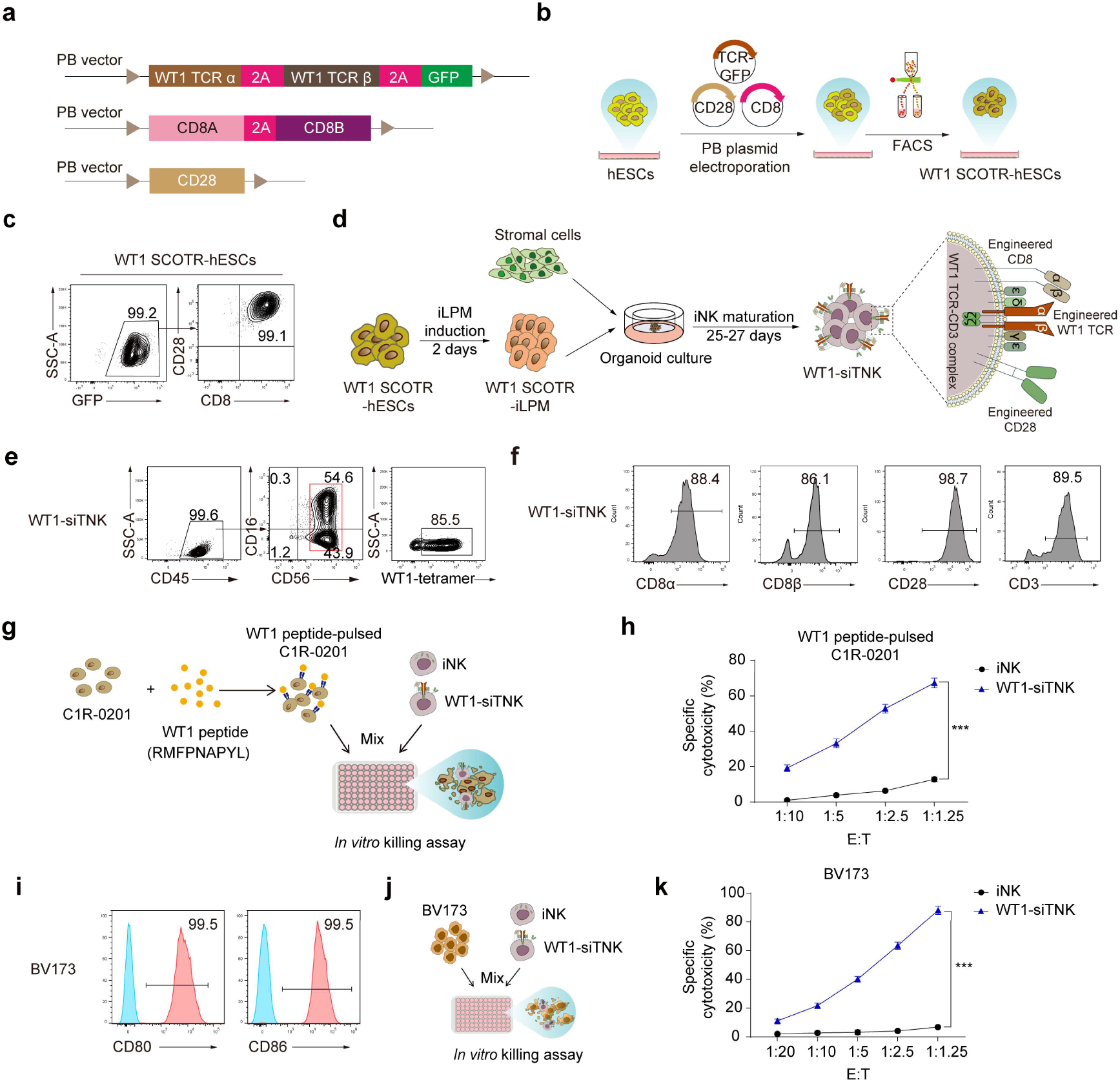
Generation and cytotoxicity evaluation of WT1-siTNK cells derived from hESCs. **a**, The design of the PB vector constructs for the WT1 TCR, CD8 coreceptor, and CD28 costimulatory molecule. **b**, Strategy diagram for the construction of WT1 SCOTR- hESCs. **c**, Flow cytometry analysis of WT1 TCR-GFP, CD8 (detected by CD8α antibody), and CD28 expression levels in WT1 SCOTR-hESCs. **d**, Schematic diagram of the induction of WT1-siTNK cells. **e**, Representative flow cytometry plots of WT1- siTNK (CD45^+^CD56^+^CD16^+/-^WT1 TCR^+^) cells. **f**, Representative histogram plots of CD8α, CD8β, CD28, and CD3 expression levels in WT1-siTNK cells. **g** and **h**, Schematic diagram (**g**) and statistical analysis (**h**) of specific cytotoxicity of iNK and WT1-siTNK cells against WT1 peptide-pulsed C1R-0201 cells (1 × 10^4^) at the indicated E:T ratios (*n* = 4 per group). **i**, The expression levels of CD80 and CD86 in engineered BV173 cells. **j** and **k**, Schematic diagram (**j**) and statistical analysis (**k**) of specific cytotoxicity of iNK and WT1-siTNK cells against BV173 cells (1 × 10^4^) at the indicated E:T ratios (*n* = 4 per group). Data were presented as mean ± SD. Statistics: two-tailed independent *t*-test (**h**, **k**), \*\*\**p* < 0.001.

### siTNK cells suppress the tumor progression in xenograft animals

To evaluate the anti-tumor efficacy of hPSC-derived siTNK cells *in vivo*, we established a human cancer xenograft mouse model by transplanting the luciferase-expressing Raji- 1101-EBV peptide (Raji-1101-EBV peptide-LUC) cells into NCG immune-deficient mice. The NCG mice were injected with Raji-1101-EBV peptide-LUC cells (1 × 10^5^ cells/mouse) on day -1 and received 1.5 Gy irradiation on day 0. Subsequently, iNK or EBV-siTNK cells were intravenously injected into the tumor-bearing animals (5 × 10^6^ cells/mouse) on day 0, day 3, and day 6, while phosphate-buffered saline (PBS) was intravenously injected as control (**Fig. 4a**). Bioluminescence imaging (BLI) revealed that EBV-siTNK cells exhibited significantly stronger tumor-killing capabilities than iNK cells (**Fig. 4b,c**). Mice treated with EBV-siTNK cells survived much longer than those treated with iNK cells or PBS (PBS: 24 days; iNK: 25 days; EBV-siTNK: 45 days; *p* < 0.001) (**Fig. 4d**). These data confirmed that EBV-siTNK cells improved the efficacy of NK cell therapy against EBV-positive tumors *in vivo*. To investigate the mechanism by which TCR enhanced iNK cell function, we performed single-cell RNA sequencing (scRNA-seq) of EBV-siTNK cells. Differential gene expression analysis indicated that 776 genes were significantly upregulated in EBV-siTNK cells relative to their expression in iNK cells (a published scRNA-seq dataset, GSA (HRA001609)). Gene ontology (GO) analysis of these 776 genes revealed that T cell-related genes were significantly enriched in EBV-siTNK cells **(Extended Data Fig. 4a)**. Further gene set enrichment analysis (GSEA) confirmed the upregulation of T-cell signaling and costimulatory pathways in EBV-siTNK cells **(Extended Data Fig. 4b,c)**. In conclusion, EBV-siTNK cells upregulated T cell-related gene expression to improve the specific cytotoxicity against target cells.

**Fig. 4.**
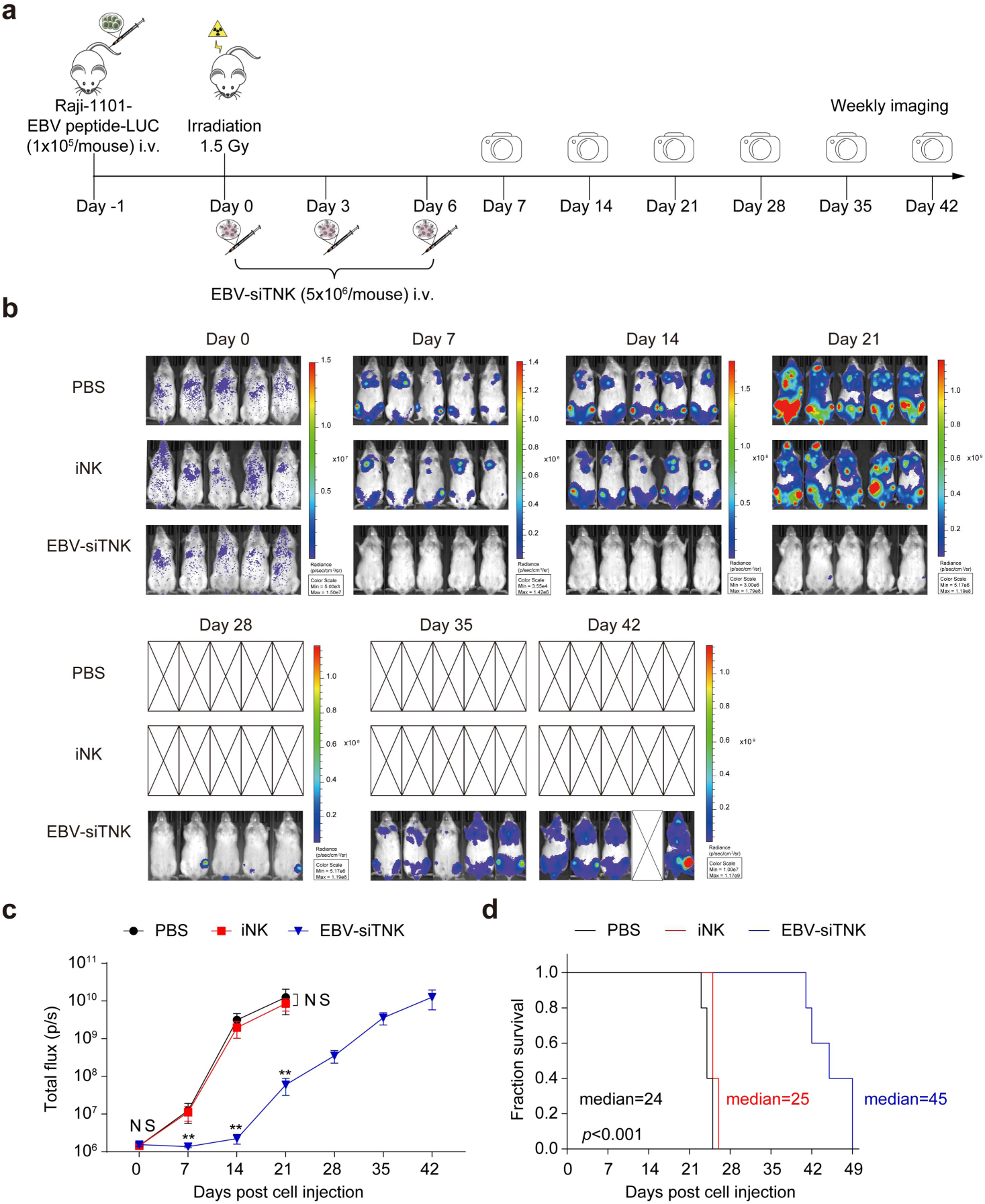
EBV-siTNK cells exhibit robust efficacy in suppressing tumor progression *in vivo*. **a**, Schematic diagram of evaluating the cytotoxicity of EBV-siTNK cells *in vivo*. NCG mice were intravenously injected with the Raji-1101-EBV peptide-LUC cells (1 × 10^5^ cells/mouse) to construct the xenograft models on Day -1. These models were intravenously injected with the iNK or EBV-siTNK cells (5 × 10^6^ cells/mouse) on Day 0, Day 3, and Day 6, respectively. BLI was performed weekly to assess the tumor burden. **b**, BLI images of xenograft models of each group (PBS, iNK cells, and EBV- siTNK cells, *n* = 5 per group). The radiance indicates tumor burden. **c**, Total flux (p/s) of the xenograft models measured at the indicated time points (*n* = 5 mice in each group). **d**, Kaplan-Meier survival curves of the xenograft models (PBS, iNK cells, and EBV- siTNK cells, *n* = 5 per group) (*p* < 0.001, Log-rank test). Median survival: PBS, 24 days; iNK, 25 days; EBV-siTNK, 45 days. Data were presented as mean ± SD. Statistics: two-tailed independent *t*-test (**c**). NS, not significant, \*\**p* < 0.01.

## Discussion

In this study, we developed a novel strategy to obtain abundant siTNK cells, which expressed an antigen-specific TCR, CD8 coreceptor, and CD28 costimulatory molecule, from engineered hPSCs. The siTNK cells integrate the inherent broad-spectrum killing activity of NK cells and the specific killing activity conferred by T cell receptors.

Through scRNA-seq of hPSC-derived iNK cells and primary NK cells, we found that iNK cells inherently express all four CD3 subunits (*CD3D*, *CD3E*, *CD3G*, and *CD247*) at the transcriptome level, while primary NK cells only express two (*CD3E* and *CD247*). This intrinsic feature enabled the efficient assembly of exogenous TCR-CD3 complex in iNK cells (72.7% vs. 9.7% in primary NK cells), suggesting that iNK cells have unique advantages compared with primary NK cells for TCR complex engineering. The SCOTR complex strategy was successfully validated in iNK cells derived from both hESCs and iPSCs. Harnessing the advantages of hPSCs, including unlimited cell sources, clonal selection for multiple genetic editing and engineering, homogeneity, and off-the-shelf features, the generation of siTNK cells from SCOTR-hPSCs successfully circumvents the issues encountered in producing TCR-NK cells from primary NK cells or NK cell lines^31,32^. Notably, siTNK cells demonstrated superior cytotoxicity compared with TCR-T cells when killing the same tumor. Some NK cell activating receptors utilize CD3ζ for signaling cascade initiation^33^. Importantly, NK cells respond to and kill target cells in a quicker manner than T cells^34,35^. Therefore, the TCR-specific activation pathway mediated by the exogenously introduced SCOTR complex can further enhance the tumor-killing capability of iNK cells.

Both EBV-siTNK cells and WT1-siTNK cells demonstrated intracellular antigen- mediated tumor-targeting and cytotoxic capabilities, proving that the SCOTR complex strategy is suitable for general tumor-associated intracellular antigen TCR assembly.

Besides, engineering TCRs into NK cells avoids TCR mispairing, a common issue observed in establishing tumor antigen-associated TCR-T cells^9–11^. Therefore, the siTNK cells offer a *de novo* tool for characterizing any TCRs, broadening their therapeutic potential.

We utilized a xenograft mouse model to evaluate the *in vivo* therapeutic effect of siTNK cells. The results demonstrated that the treatment group infused with siTNK cells (median survival time: 45 days) could significantly prolong the survival of tumor- bearing mice compared with the iNK treatment group (median survival time: 25 days). Long-term tumor remission might be achieved once the shortcoming of poor persistence *in vivo* of siTNK cells is overcome^36^. In general, approaches to prolong the persistence of NK cells include *in vivo* infusion of cytokines (e.g., IL-15)^37^, genetic modification of NK cells by blocking intracellular immune checkpoints^38,39^, and increasing the administered NK cell doses^40^. Recently, our group successfully generated CXCR4-expressing iNK progenitor cells and iNK cells from CXCR4-expressing hPSCs, and these cells exhibited promising therapeutic effects in tumor models^41,42^. CXCR4 enhanced the homing ability of hPSC-derived iNK cells and prolonged their persistence within the body. These findings offer valuable insights for the further enhancement of the functions of siTNK cells.

In conclusion, siTNK cells provide a brand-new cell resource for cancer treatment with significant public health implications.

## Statistical analysis

All quantitative analyses were performed using SPSS (IBM SPSS Statistics 25). The Shapiro-Wilk test was used to evaluate the normal distribution of the data. Quantitative differences (mean ± SD) between samples were compared using two-tailed independent *t*-test and Mann-Whitney U test. The *p* values were two-sided, and *p* < 0.05 was considered statistically significant. Survival curves for tumor-bearing models were plotted using the Kaplan-Meier method and compared between groups using the log-rank (Mantel-Cox) test. Statistical analysis was performed using GraphPad Prism 9 software.

## Data availability

The scRNA-seq data have been uploaded to the Genome Sequence Archive public database (HRA010093). R scripts with R files used to analyze scRNA-seq data have been deposited on Github and are publicly available (https://github.com/lyqbiocc/scRNAseq-of-sCOTR-iNK). Other relevant information or data is available from the corresponding authors upon reasonable request.

## Acknowledgments

This work was supported by grants from the National Key R&D Program of China (2024YFA1108302), the National Natural Science Foundation of China (82450001, 81925002, 82300132, and 82470120), and the Noncommunicable Chronic Diseases-National Science and Technology Major Project (No. 2023ZD0501200).

## Author contributions

Conceptualization: J.Y.W., M.Y.Z.; Methodology: X.J.Z., F.Z., F.X.H., P.C.L., Z.Q.W., Y.H.L., Z.Y.X., Y.W., L.Q.Z., H.M.Q., Y.Y.S., Y.C., J.X.W., J.C.X., Y.Q.Z.; Investigation: X.J.Z, F.Z.; Visualization: X.J.Z., F.Z., Q.T.W., Y.Q.L, C.Y.Z.; Funding acquisition: J.Y.W., M.Y.Z.; Project administration: T.J.W., Y.P.Z., L.J.L.; Supervision: J.Y.W., M.Y.Z., C.X.X., D.H.H., T.J.W., F.X.H.; Writing – original draft: X.J.Z., C.X.X., D.H.H., M.Y.Z.; Writing – review & editing: X.J.Z., J.Y.W., M.Y.Z.

---

**Extended Data Fig. 1.**
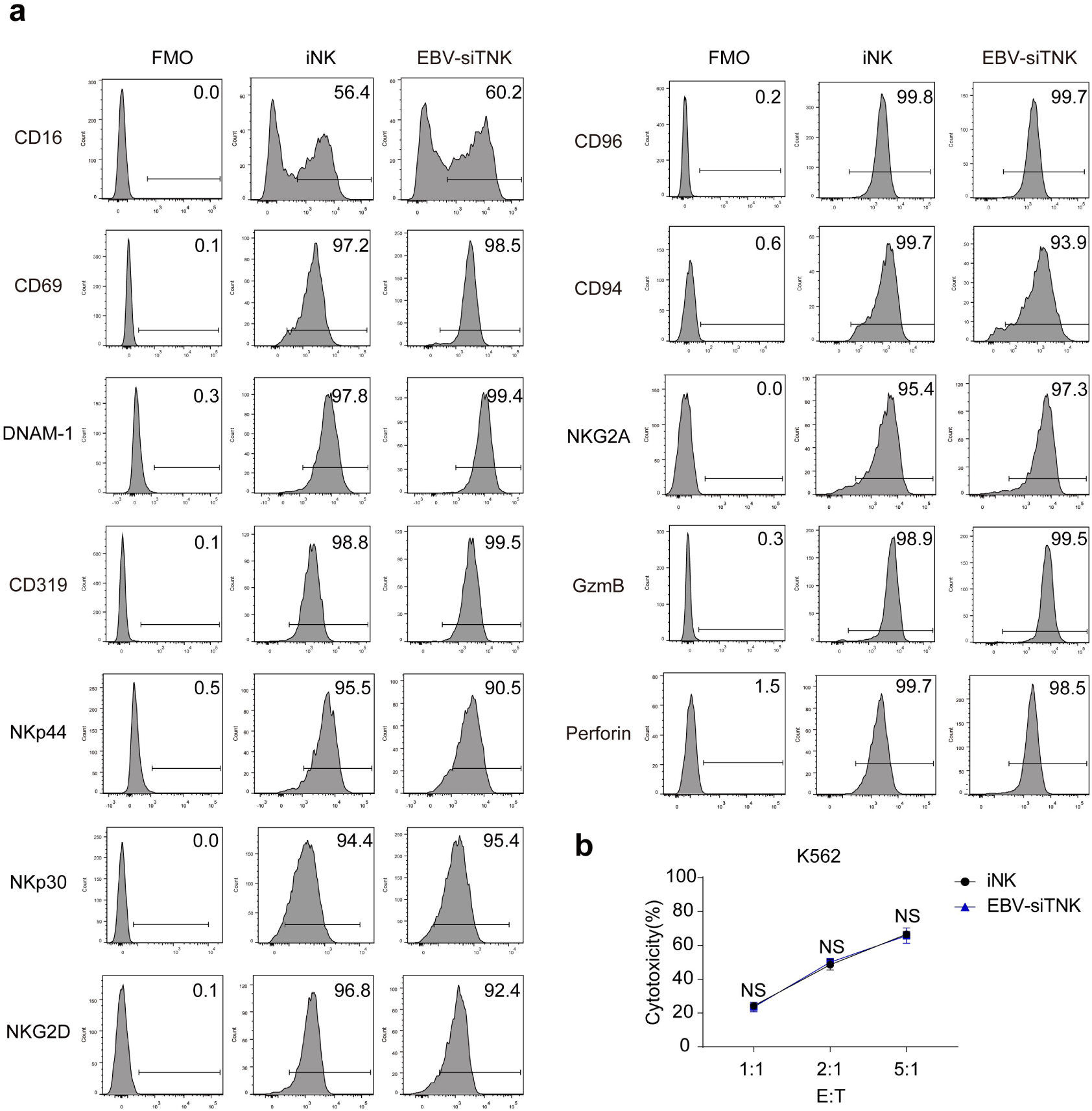
Molecular features and non-specific cytotoxicity of EBV- siTNK cells. **a**, Flow cytometry analysis of the expression levels of NK cell receptors and effectors (CD16, CD69, DNAM-1, CD319, NKp44, NKp30, NKG2D, CD96, CD94, NKG2A, GzmB, and Perforin). **b**, Cytotoxicity analysis of iNK and EBV-siTNK cells against K562 tumor cells at the indicated effector to target (E:T) ratios after 4 h incubation (*n* = 3 per group). Data are represented as mean ± SD. Statistics: two-tailed independent *t*-test. NS, not significant.

**Extended Data Fig. 2.**
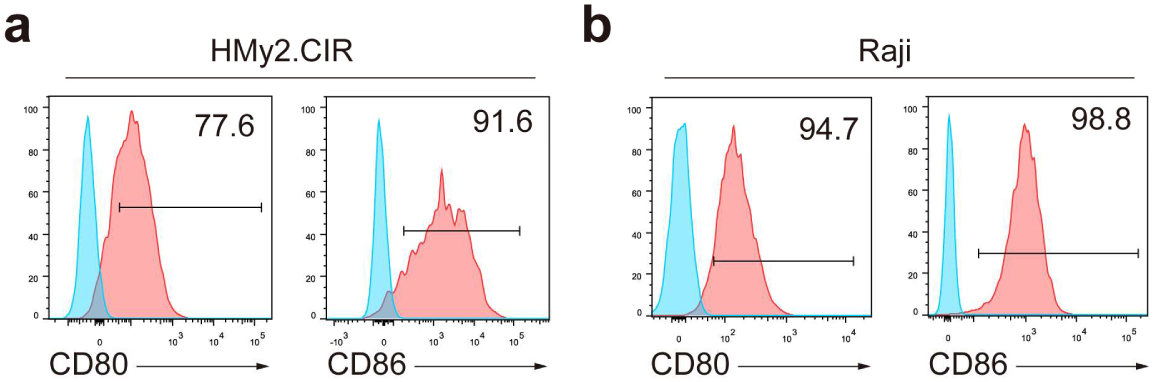
The expression levels of CD80 and CD86 in HMy2.CIR and Raji cells. **a** and **b**, Representative histogram plots of CD80 and CD86 in HMy2.CIR (**a**) and Raji (**b**) cells.

**Extended Data Fig. 3.**
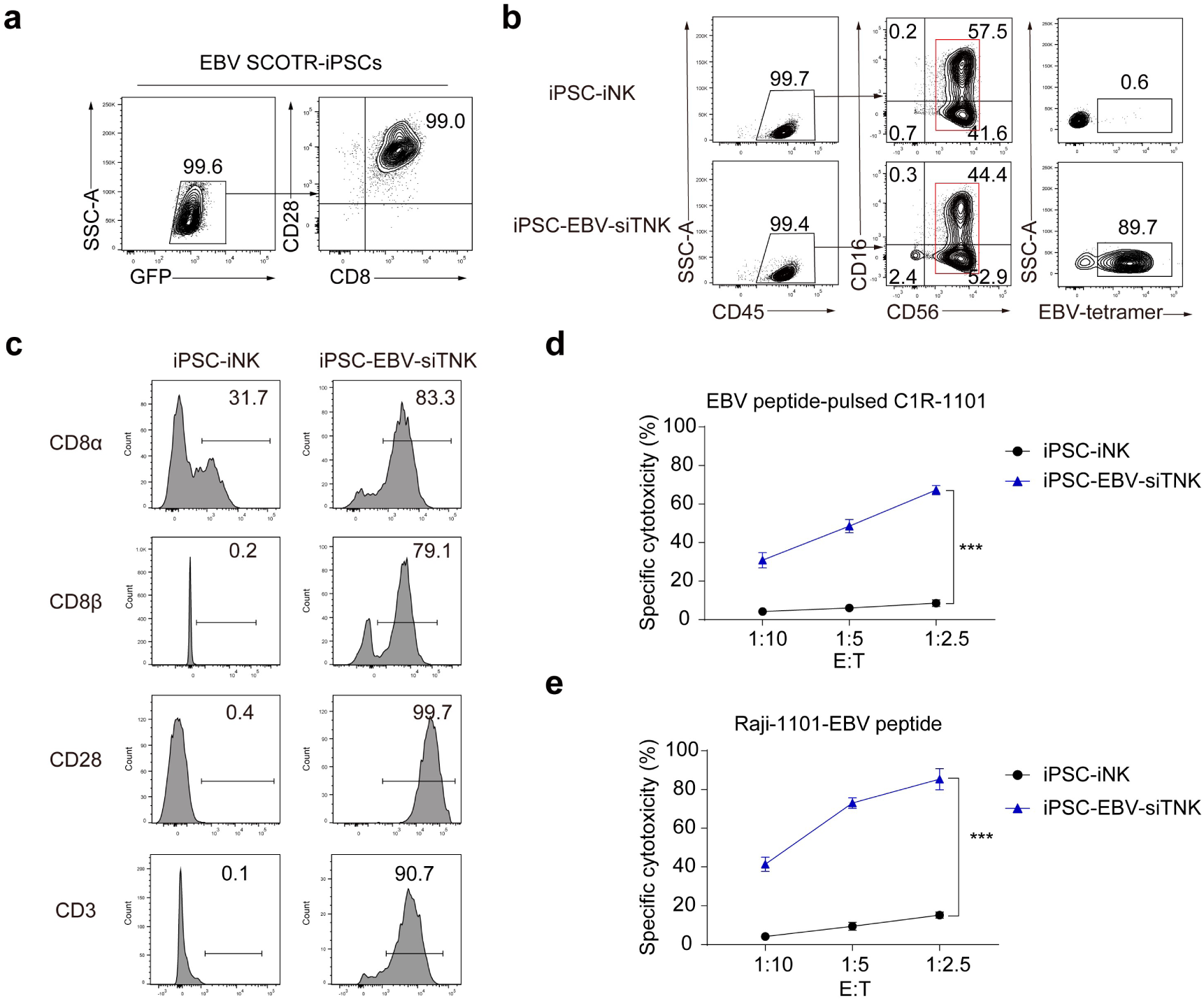
Generation and cytotoxicity evaluation of EBV-siTNK cells derived from iPSCs. **a**, Flow cytometry analysis of EBV TCR-GFP, CD8 (detected by CD8α antibody), and CD28 expression levels in EBV SCOTR-iPSCs. **b**, Representative flow cytometry plots of iPSC-iNK and iPSC-EBV-siTNK (CD45^+^CD56^+^CD16^+/-^EBV TCR^+^) cells. **c**, Representative histogram plots of CD8α, CD8β, CD28, and CD3 (detected by CD3ε antibody) expression levels in iPSC-iNK and iPSC-EBV-siTNK cells. **d** and **e**, Statistics of specific cytotoxicity analysis of iPSC-iNK and iPSC-EBV-siTNK cells against EBV peptide-pulsed C1R-1101 cells (**d**) and Raji-1101-EBV peptide cells (**e**) at the indicated E:T ratios (*n* = 4 per group). Statistics: two-tailed independent *t*-test, \*\*\**p* < 0.001.

**Extended Data Fig. 4.**
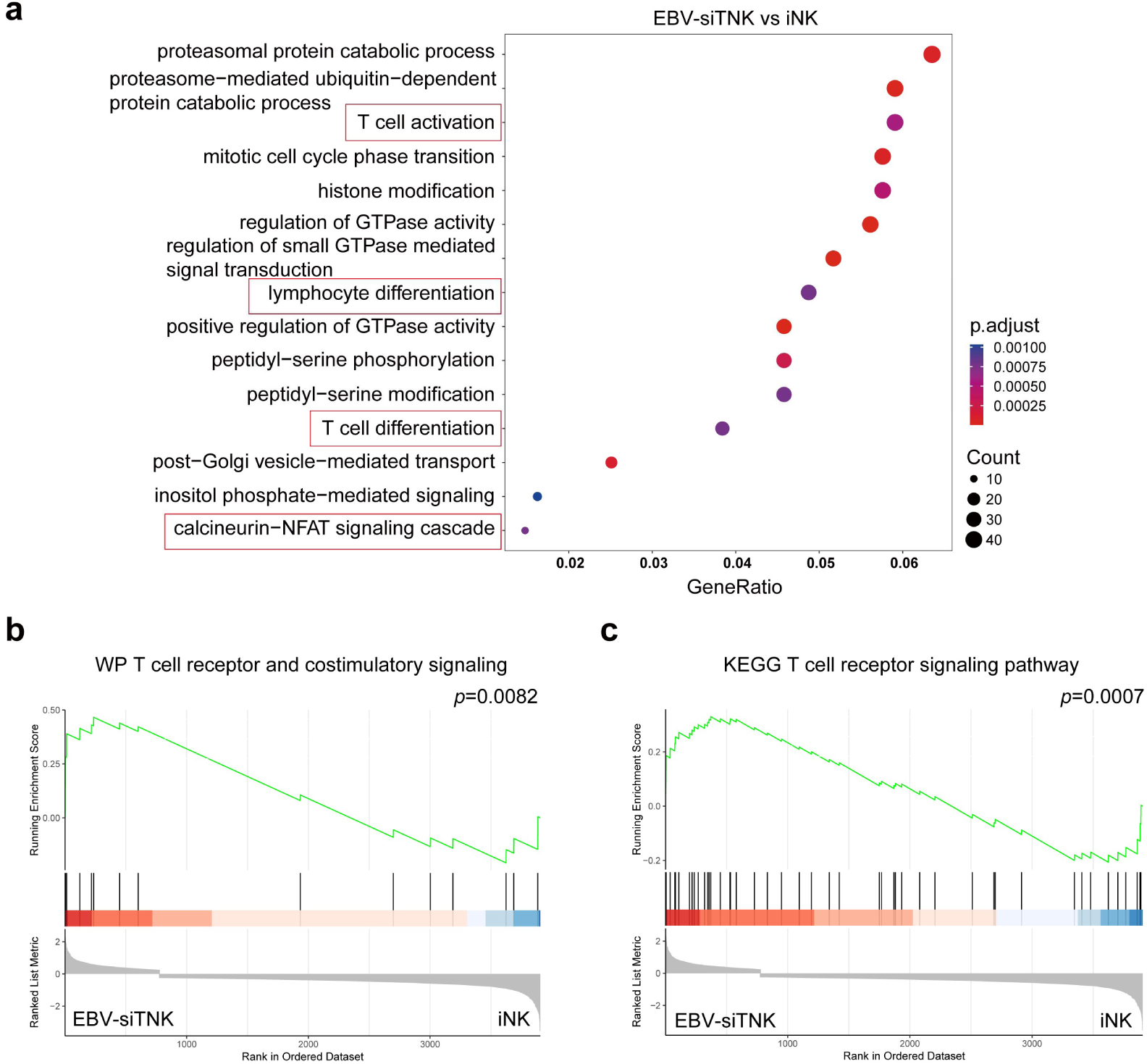
Transcriptomic analysis of TCR signaling associated pathways in EBV-siTNK cells. **a**, GO enrichment analysis of the 776 up-regulated differentially expressed genes between EBV-siTNK and iNK cells. Each symbol represents a GO term (noted in the plot), color indicates adjusted *p* value (p.adjust (significance of the GO term)), and symbol size is proportional to the number of genes. The differentially expressed genes were identified by the FindMarkers function of the Seurat package with min.pct = 0.25 and logfc.threshold = 0.25. The GO enrichment analysis was performed with the clusterProfiler package. The raw data of iNK cells were downloaded from GSA (HRA001609). **b**, GSEA of genes in WP T cell receptor and costimulatory signaling. The gene set was obtained from MSigDB. The clusterProfiler package was used for GSEA. **c**, GSEA of genes in KEGG T cell receptor signaling pathway. The gene set was obtained from MSigDB. The clusterProfiler package was used for GSEA.

